# Treating symptomatic infections and the co-evolution of virulence and drug resistance

**DOI:** 10.1101/2020.02.29.970905

**Authors:** Samuel Alizon

## Abstract

Antimicrobial therapeutic treatments are by definition applied after the onset of symptoms, which tend to correlate with infection severity. Using mathematical epidemiology models, I explore how this link affects the coevolutionary dynamics between the virulence of an infection, measured via host mortality rate, and its susceptibility to chemotherapy. I show that unless resistance pre-exists in the population, drug-resistant infections are initially more virulent than drug-sensitive ones. As the epidemic unfolds, virulence is more counter-selected in drug-sensitive than in drug-resistant infections. This difference decreases over time and, eventually, the exact shape of genetic trade-offs govern long-term evolutionary dynamics. Using adaptive dynamics, I show that two types of evolutionary stable strategies (ESS) may be reached in the context of this simple model and that, depending on the parameter values, an ESS may only be locally stable. In general, the more the treatment rate increases with virulence, the lower the ESS value. Overall, both on the short-term and long-term, having treatment rate depend on infection virulence tend to favour less virulent strains in drug-sensitive infections. These results highlight the importance of the feedbacks between epidemiology, public health policies and parasite evolution, and have implications for the monitoring of virulence evolution.

## Introduction

Chemotherapies represent a major selective force for microbes. It is now commonly accepted that the administration of these treatments always eventually selects for resistant strains (AF Read, Day, et al., 2011). However, drug resistance is not the only infection life-history trait that may evolve, and mathematical (Gandon, Mackinnon, et al., 2001; Gandon and Michalakis, 2000) and some animal (AF Read, Baigent, et al., 2015; Schneider et al., 2012) models support the idea that treatments can affect the evolution of virulence, defined here as the decrease in host fitness due to the infection (A Read, 1994).

This study analyses the evolution of antimicrobial resistance in parasite strains that can vary in virulence. This trait was long seen as binary, following Pasteur’s view was that ‘virulent strains killed whereas attenuated strains did not’ (Mendelsohn, 2002), which led to the successful identification of ‘virulence factors’, some of which can be associated with drug resistance (Alcalde-Rico et al., 2016; Beceiro et al., 2013; Copin et al., 2019; Geisinger et al., 2018; Giraud et al., 2017; Guillard et al., 2016; Van Tyne and Gilmore, 2014). However, the field is increasingly considering virulence as a quantitative trait under the control of many genes (Casadevall et al., 2011) but experimental data is still more limited.

In epidemiology, there is a rich literature on the evolution and spread of drug resistance (Spicknall et al., 2013). Some studies also investigate the coevolution between two drug-resistance loci (Day and Gandon, 2012). However, the coevolution of drug resistance and virulence is rarely studied. The problem may appear simplistic at first because drug-resistant infections being more difficult to treat, their virulence is likely is higher than drug-sensitive ones in treated hosts. But coevolution can be less straightforward. For instance, as documented for several bacterial species, there can be direct genetic associations between the two traits due to genes with pleiotropic action, which means they can affect multiple infection life-history traits for reviews, see Beceiro et al., 2013; Guillard et al., 2016. Note that in these studies, virulence is usually a binary rather than a continuous trait. A final link between these two traits, which is the focus of this study, is created by public health policies since less virulent strains are less likely to be treated than more virulent ones. Symptoms are, almost by definition, the driver of curative treatments (Canguilhem, 1978). The relationship between virulence, symptoms and treatment rate is difficult to quantify, even if it is the basis of many discussions the medical field (Ewald, 1980; Porco et al., 2005; Tuckett, 2013). In some cases, the link is clear. For example, before the advent of the test-and-treat policy, international guidelines recommended initiating antiretroviral therapy in HIV infected patients only when CD4T-cell counts fell below a given threshold (Hammer et al., 2006), an event known to be associated with the virulence of the infecting strain (Fraser, Lythgoe, et al., 2014). In the case of malaria, some advocate for the treatment of asymptomatic infections (Chen et al., 2016). Finally, this issue is particularly timely in the context of overtreatment and the evolution of antimicrobial resistance (Lin et al., 2012).

Classical theory predicts that if virulence is adaptive for the parasite, defined here to encompass both micro-and macro-parasites, host qualitative resistance selects for increased levels of virulence (Gandon and Michalakis, 2000). If host resistance is quantitative, which would better correspond to efficient drug treatments, virulence is not affected in simple models (Alizon and van Baalen, 2005). However, if multiple infections are allowed in the system, their prevalence decreases, potentially affecting virulence evolution (Gandon, Mackinnon, et al., 2001). Although the long term effect of host resistance (or prophylactic treatments) on virulence evolution may be equivalent to therapeutic treatments in simple models, on the short term, however, the fact that only a fraction of hosts are treated based on infection life-history traits may influence evolutionary dynamics. Furthermore, the nature of the tradeoffs between infection life-history traits can also affects the effect of treatment rate on the evolutionarily stable level of virulence (Porco et al., 2005).

Variations in infection life-history traits may occur in different ways. At one extreme, drug resistance can evolve rapidly because one or a few mutations can have large phenotypic effects; as illustrated for instance in the case of *Mycobacterium tuberculosis* (Telenti, 1997), *Plasmodium falciparum* (White, 2004) or influenza virus (Foll et al., 2014). Conversely, quantitative traits such as virulence or transmission rate tend to evolve on a rougher fitness landscape, as estimated for instance in the case of Human Immunodeficiency Virus (HIV) infections through the virus load proxy (Hinkley et al., 2011). Furthermore, drug-resistance is known to be associated with fitness costs for the parasite that can lead to decreased transmission, as shown for bacteria (Andersson and Hughes, 2010; Luciani et al., 2009) but also viruses (Kühnert et al., 2018).

Here, I adopt an evolutionary epidemiology standpoint and introduce quantitative strain variations in virulence as well as qualitative variations in infection drug resistance. The model is generic and can be re-scaled in time to capture either acute, influenza-like, infections with high transmission rate or chronic, HIV-like, infections with lower transmission rate. A key assumption is that the rate at which an infected host receives treatment can correlate with infection virulence. The hypothesis tested here is that this correlation can generate coevolutionary dynamics between virulence and drug resistance on the short and long terms. These are explored by applying the Price equation (Day and Proulx, 2004) and the adaptative dynamics (Dieckmann, 2002) frameworks, respectively, to study virulence evolution.

## Model and Methods

### The SI model

I assume a classical Susceptible-Infected model (Keeling and Rohani, 2008) in which infections caused by a strain *i* are either drug-sensitive or drug-resistant (Figure 1). Let us denote by *S* the density of susceptible hosts, by *I_i_* that of drug-sensitive infections by strain *i* and by 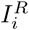 that of drug-resistant infections by strain *i*. Infection traits are assumed to be ‘heritable’ such that the infection of a susceptible host by a drug-sensitive infection by strain *i* (at a rate *β_i_*) yields a drug-sensitive infection by strain *i*. New drug-resistant infections appear in two ways: either through the infection of a susceptible host by a drug-resistant infection (at a rate 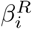), or following the treatment failure of a drug-sensitive host. The latter event occurs at a rate *ρ_i_γ_i_*, where *γ_i_* is the treatment rate and *ρ_i_* is the proportion of drug failure. For simplicity, hosts are assumed not to recover from drug-resistant infections. Finally, new hosts enter the system at a density-independent rate λ and die with a natural mortality rate *μ.* Infected hosts experience additional mortality, i.e. virulence, *α_R_* for drug-sensitive infections and *α_R_* for drug-resistant infections. Note that in the model, virulence and transmission rate can differ in drug-sensitive and drug-resistant infections. Biologically, these trait differences could originate from fitness-costs associated with drug resistance (Andersson and Hughes, 2010). Finally, for infected hosts, I assume that they can also recover naturally from the infection (i.e. independently of the treatment rate *γ_i_*) at a constant rate *ν*. Individuals who die or recover naturally are removed from the system, which implicitly assumes lifelong immunity.

**Figure 1.**
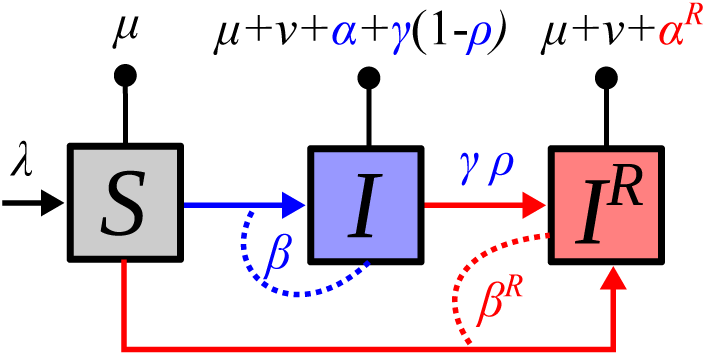
Epidemiological model flow diagram. Drug-resistant infections (in red) can originate from direct transmission or treatment failure in drug-sensitive infections (in blue). Dashed lines show infection events, arrows transition between states and lines with a circle death or recovery events. Individuals who die or recover from the infection are removed from the system. Parameter notations are detailed in Table 1.

**Table 1.** Model parameters and notations used.

| Notation | Description |
| --- | --- |
| $\alpha$ | virulence |
| $\beta$ | transmission rate |
| $\gamma$ | treatment rate |
| $\rho$ | treatment failure ratio |
| $\lambda$ | host input rate |
| $\mu$ | host baseline mortality rate |
| $\nu$ | natural recovery rate |
| $i$ | relative to parasite strain $i$ |
| $m$ | relative to a mutant strain |
| $R$ | relative to drug-resistant infections |
| $*$ | relative to an evolutionary stable strategy (ESS) |
| $n$ | number of parasite strains |
| $p_i$ | proportion of strain $i$ in drug-sensitive infections |
| $q_i$ | proportion of strain $i$ in drug-resistant infections |
| $\bar{x}^A$ | average value of trait $x$ in compartment $A$ (drug-resistant or sensitive) |
| $\text{Cov}^A(x, y)$ | genetic covariance between traits $x$ and $y$ measured in compartment $A$ |
| $\text{Var}^A(x)$ | genetic variance in traits $x$ measured in compartment $A$ |
| $\phi$ | invasion fitness |
| $a$ | variation in virulence in drug-resistant infections |
| $b$ | variation in transmission rate in drug-resistant infections |
| $p$ | transmission-virulence trade-off shape parameter |
| $q$ | treatment-virulence trade-off shape parameter |

The dynamics of the system are captured by the following set of Ordinary Differential Equations (ODEs):

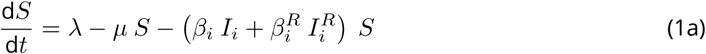

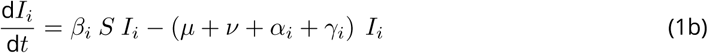

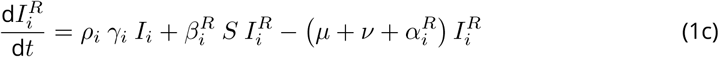

For *α, β, γ* and *ρ*, variations between infections are assumed to depend on genetic differences between strains and to vary slowly. Conversely, drug resistance is assumed to be a more labile trait. A possible biological interpretation is that it reflects the state of the majority of the parasites in an infected host at a given time point. Let us allow this trait to vary, while the others remain constant; whence the existence of the two classes *I_i_* and 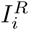 and the possibility to ‘mutate’ from the former to the latter (reversions are neglected in this model). Genetic mutations from a strain *i* to a strain *j* are neglected because if the mutation kernel is symmetric these terms cancel out (Day and Proulx, 2004).

For simplicity, I assume that 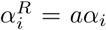 and 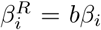, meaning that all strains undergo similar variations in infection traits with the acquisition of drug-resistance. Note that this assumption implies that 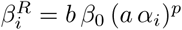. Typically, if drug resistance is costly for the parasite, we expect to have *a* > 1 (i.e. increased virulence) or 0 ≤ b < 1 (decreased transmission rate), where a and 1/b can be interpreted as fitness costs.

### Short-term evolutionary dynamics

To follow the trait dynamics at the population level, I introduce 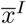(resp. 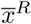) as the average trait value of *x* in the drug-sensitive (resp. drug-resistant) compartment. Mathematically, 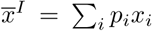, where *p_i_ = I_i_/I_T_* is the fraction of strain *i* among all the drug-sensitive infections. Similarly, 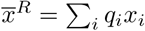, where 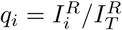 is the fraction of strain *i* among all the drug-resistant infections.

The dynamics of the total densities of infected hosts (*I*_*T*_ and 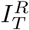) are governed by the following ODEs:

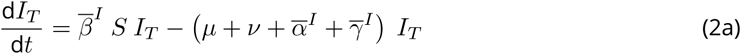

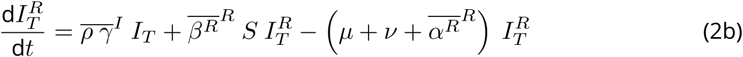

Even for this simple SI model, each strain i is characterised by 6 infection traits (*α*_*i*_,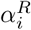,*β*_*i*_,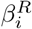,*γ*_*i*_ and *ρ*_*i*_). Furthermore, at the population level, the average values of each of these traits can differ in the drug-sensitive and drug-resistant compartments (*I* and *I^R^*). Note that there is an identifiability issue between the treatment failure ratio (*ρ*) and the treatment rate (*γ*) since both are averaged simultaneously in equation 2b. Note also that, in this simple system, natural mortality and recovery rates (*μ* and *ν*) have the same effect on the dynamics, which means either can be set to 0 without affecting short-term dynamics.

After some calculations using the Price equation formalism introduced by Day and Proulx (2004) and described in Appendix S1, I obtain the following ODEs to capture the dynamics of the average trait value of a trait *x* in the drug-sensitive compartment and a trait *y* in the drug-resistant compartment:

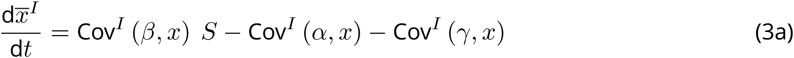

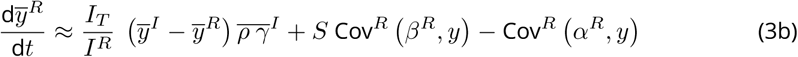

where Cov*^A^* (*x*, *y*) is the genetic covariance between traits *x* and *y* measured in compartment *A*. This will be discussed further in the Results but, for example, a positive covariance between treatment rate and virulence Cov*^I^* (*γ*, *α*) > 0 means that more virulent infections are treated more.

### Long-term evolution

System 1 accepts three equilibria for 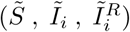, which are detailed in Appendix S1. The first equilibrium corresponds to a disease-free equilibrium, where the parasite goes extinct (λ/*μ*, 0,0). In the second, resistant infections are only maintained via treatment failure (*γ_i_ ρ_i_*). Without it, the equilibrium simplifies into 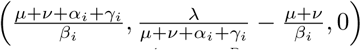. In the third equilibrium, all infections are found to be drug-resistant: 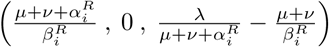.

By adopting an adaptive dynamics approach, it is possible to investigate the convergence and evolutionary stability of fixed points (Dieckmann, 2002). Practically, we study a version of system 1b for 2 strains, where one of them, referred to as the ‘resident’ and denoted by the subscript *r*, is assumed to be at its endemic equilibrium, and the other, the ‘mutant’ denoted by the subscript *m*, is assumed to be rare.

As described in Appendix S1, the next generation theorem (Diekmann et al., 1990; Driessche and Watmough, 2002; Hurford et al., 2010) tells us that the fate of the mutant is governed by the largest of the two following eigenvalues:

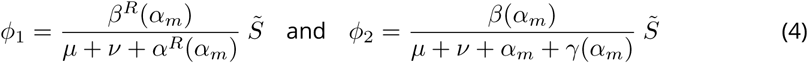

where 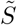 is the equilibrium density of susceptible hosts in a ‘resident’ population before the emergence of a mutant strain with virulence *α_m_*. Note that transmission and treatment rates, as well as the virulence of drug-resistant infections, can here be functions of virulence in drug-sensitive infections. As in the short-term evolution model, we can, for example, model the fact that more virulent infections are treated more by assuming that the derivative of *γ*(*α*) with respect to α is positive.

If *φ* = max(*φ*_1_,*φ*_2_) > 1, the mutant invades the resident system and becomes the new resident.

One possible biological interpretation of the two eigenvalues *φ*_1_ and *φ*_2_ is that either the parasite spreads via drug-resistant or drug-sensitive infections. This dichotomy originates from the assumption that the mutant is initially rare, which makes the generation of drug-resistant infections via mutation a second-order term that vanishes when evaluating the eigenvalues of the Jacobian matrix.

Classically, a virulence level will be defined as evolutionary stable and denoted *α*\* if it is such that

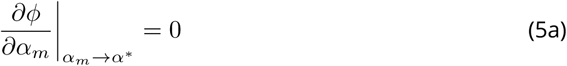

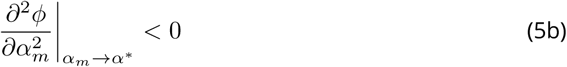

### Numerical simulations

To illustrate the analytical results, I perform numerical simulations with *n* = 20 strains the traits of which are drawn at random (the `R` script used with the default parameter values is in Appendix S2). These simulations assume a positive covariance between virulence (*α_i_*) and treatment rates (*γ_i_*), therefore considering that more virulent infections are treated more. Practically, I assume that the values of *α_i_*, and *γ_i_*, are drawn in a multivariate normal distribution such that Var(*α*) = 1/6, Var(*γ*) = 1/12, and Cov(*α, γ*) = 1/10. The resulting (*α_i_*, *γ_i_*) pairs are shown in Figure S3A.

The default time unit in the simulations is weeks. However, since the demography plays a very minor role in the model (it mainly determines the initial number of susceptible hosts), the system can be rescaled to any time unit without affecting the results qualitatively.

Following empirical and experimental data (Anderson and May, 1982; de Roode et al., 2008; Doumayrou et al., 2012; Dwyer et al., 1990; Fraser, Hollingsworth, et al., 2007; Råberg, 2012; Williams et al., 2014), the transmission rate is assumed to be traded-off against infection duration due to the cost virulence such that

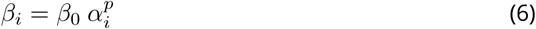

where *β*_0_ is a scaling constant. If 0 < *p <* 1, theory predicts there is an intermediate evolutionary stable level of virulence (Alizon and van Baalen, 2005; van Baalen and Sabelis, 1995).

In the Appendix, I also investigate the effect of noise in the trade-off relationship. Practically, this is modelled by assuming that the realised transmission rate is normally distributed around the value predicted from the trade-off, i.e. with mean 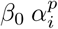 and variance 0.2 (see Figure S3B).

Finally, for simplicity, I assume in the simulations that drug-resistance infections are similar to drug-sensitive infections (*a* = *b* = 1).

## Results

### Virulence rapid evolution

The Price equation formalism gives us a qualitative understanding of the selective forces acting on infection traits. The short term dynamics of our main trait of interest, virulence in drug-sensitive infections, are obtained from equation system 3:

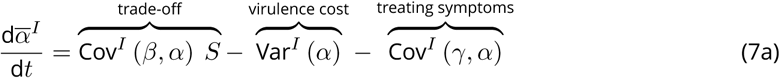

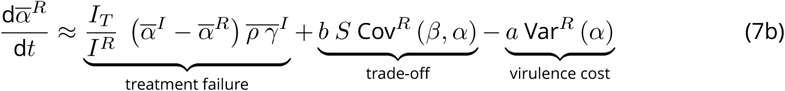

where *a* ≥ 1 and 0 < b ≤ 1 are parameters that relate to the fitness cost associated with drug-resistance.

In drug-sensitive infections, virulence evolutionary dynamics are similar to that described for classical SIR systems (Day and Proulx, 2004). The first term in the right-hand side of equation 7a indicates that the average virulence increases (i.e. more virulent strains spread more) if there is a positive covariance between transmission rate and virulence, which is also known as the transmission-virulence trade-off hypothesis (Alizon and Michalakis, 2015). This effect is amplified by the availability of susceptible hosts (*S*), as demonstrated experimentally in a bacteria-phage system (Berngruber et al., 2013). According to the second term, virulence is costly and if there is genetic variation for this trait, more virulent strains are counter-selected; a phenomenon introduced more than a century ago by Theobald Smith as the ‘law of declining virulence’ (Méthot, 2012).

The third term in equation 7a deserves some explanations because it involves the rate at which infections are treated (*γ_i_*). In most epidemiological models, this rate is assumed to be the same for all the strains. Here, this term allows us to assume that treatments are triggered by infection symptoms, which are themselves correlated to virulence. If the covariance is positive, the term acts as a selective force against virulence in the drug-sensitive compartment.

Virulence evolutionary dynamics in the drug-resistant infections compartment are partly governed by similar forces than in the drug-sensitive compartment. The second term in the right-hand side of equation 7b is the transmission-virulence trade-off, while the third is the law of declining virulence. The proportionality constants *a* and *b* originate from our simplifying assumption regarding the link between traits expressed both in drug-sensitive and drug-resistant infections (see the Model section).

The first term in equation 7b corresponds to treatment failure events and homogenises trait values between drug-sensitive and drug-resistant compartments. If virulence is the same in the two compartments, this term is zero. As mentioned above, variations in the fraction of treatments that fail (*ρ_i_*) cannot be separated from variations in the treatment rate (*γ_i_*). Finally, this term is weighted by the density of drug-sensitive and drug-resistant infections. Initially, infections are drug-sensitive and this treatment failure term governs most of the dynamics. It is only in a second phase that transmitted drug-resistance can matter and that the second and third terms weight in. This term is not present in 7a because reversions from drug-resistant to drug-sensitive infections are neglected in this model.

Overall, virulence in the drug-resistant compartment should be less selected against than in the drug-sensitive compartment because of the absence of covariance term between *α* and *γ* in equation 7b.

### Numerical simulations

#### Short-term evolution

To illustrate these analytical results, I perform numerical simulations with *n* = 20 strains with sets of traits drawn at random and assuming a transmission-virulence trade-off. Figure 2 shows the short term evolutionary dynamics predicted by the Price equation formalism (dashed lines) and those obtained via simulations (plain lines). For simplicity, I assume that drug-resistance infections do not incur a fitness cost.

**Figure 2.**
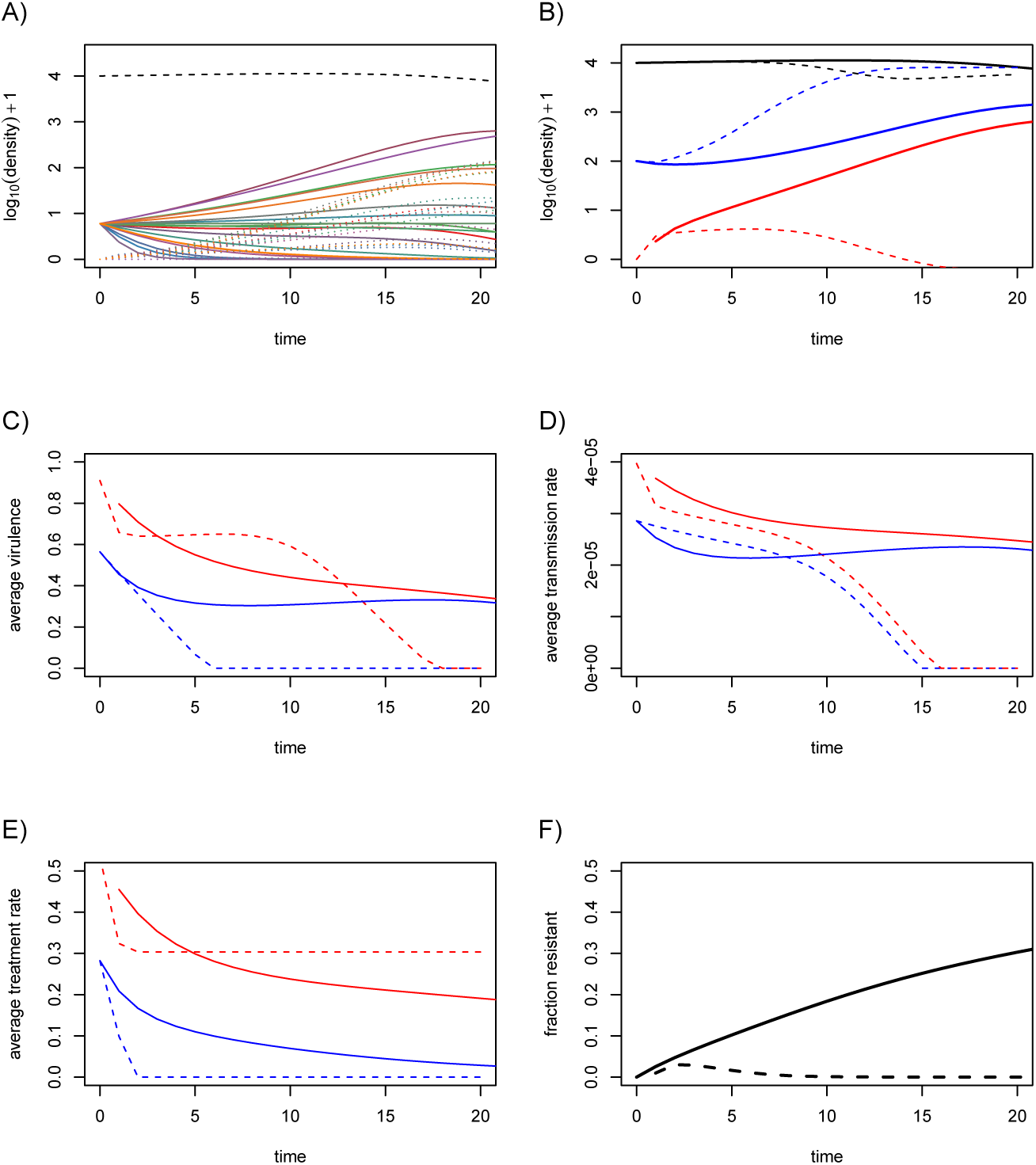
Short term evolutionary dynamics. A) Dynamics of the densities of susceptible hosts (dashed black line), of drug-susceptible infections (plain coloured lines) and drug-resistant infections (dotted coloured line). Each colour corresponds to one of the *n* =20 strains. B) Same as A but the blue line shows the total density of drug-susceptible infections and the red line the total density of drug-resistant infections. The dashed lines are the predictions from the Price equation system. C) Average virulence in the drug-sensitive (blue) and drug-resistant infections (red) for numerical multi-strain simulations (plain line) and the Price equation system (dashed line). D) Same as panel C for transmission rate. E) Same as panel C for treatment rate. F) Fraction of the infections that are drug-resistant in the multi-strain simulation (plain line) or using the Price equation (dashed line). We assume no fitness cost (*a* = *b* =1) and a transmission-virulence trade-off 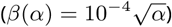. Other parameter values are λ = 0.02, *μ* = 4.5 × 10^−5^, *ν* = 0, *a* = *b* = 1, *ρ* = 0.1, Cov(*α*, *γ*) = 0.1, Var(*γ*) = 1/12, Var(*α*) = 1/6, 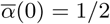, 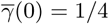, *S*(0) = 10^4^, *I_i_* = 5, and 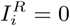. The default time unit is the week (but it can be rescaled without affecting the results qualitatively).

Figure 2A illustrates the diversity of the epidemiological dynamics among all of the 20 strains. We see that not all the strains spread during the epidemics. We also see that initially, all the infections are drug-sensitive (dotted lines start from 0). Figure 2B shows the total densities of each type of infection (drug-sensitive or drug-resistant) and the predictions from the Price equation formalism (dashed lines). These predictions are correct at first but they rapidly overestimate the spread of drug-sensitive infections (in blue) and underestimate the spread of drug-resistant infections (in red).

Focusing on average infection trait values, Figure 2C shows that, as predicted, virulence decreases faster in the drug-sensitive infections than in the drug-resistant infections. However, it also shows the limits of the Price equation because parasites eventually evolve to complete avirulence. Note that in the Price equation approach, I assumed different initial trait values for the drug-sensitive and drug-resistant compartments. Indeed, drug-resistance is initially absent in the simulations (the plain red line starts with a lag) and average infection virulence is much higher in this compartment. This is because resistant infections first emerge from treatment failure and not all strains have the same exposition to treatments. Therefore, instead of using the average virulence of all the strains in the population as our initial value, we weight these virulences by the recovery rates. This approximation is consistent with the simulated dynamics (Figure 2C).

Importantly, the mismatch in virulence observed in drug-resistant infections is not solely due to this initial effect of more virulent infections being more treated. Indeed, as shown in Supplementary Figure S1, even if initial densities of drug-sensitive and drug-resistant infections are assumed to be equal, therefore implying a pre-existence of drug resistance, the initial virulence in the two compartments are identical but there is still a more rapid decrease in virulence in the drug-sensitive compartment, as predicted by the Price equation model. This cannot be explained by a fitness difference between the two strains because there is no fitness cost in these simulations.

Because of the genetic variances and covariances, other traits coevolve with virulence. In Figure 2D, we see that the Price equation does not predict any difference in transmission rate between sensitive and resistant infections. In the simulations, the transmission rate in the drug-resistant compartment remains higher than that in the drug-sensitive compartment but this is due to the initial increase in virulence in this compartment (explained above). For the rate at which infections are treated (Figure 2E), we see a similar pattern than for virulence, which is consistent with the fact that the two are strongly correlated. Finally, Figure 2F shows the fraction of drug-resistant infections. Again, the Price equation is initially accurate but it rapidly underestimates the spread of drug resistance. From a biological standpoint, it is interesting to notice that the decrease in the average level of virulence goes along with a decrease in average treatment rate.

#### Long-term evolution

Long-term simulations show that the parasite can persist (Figure 3A). We also see that the average virulence in the drug-sensitive compartment (in blue) increases before reaching avirulence (Figure 3B). After a long transient phase, it even drops below the average virulence in the drug-resistant compartment (in red). Figure 3C shows that strains can be maintained by different types of infections. For instance, for times between 100 and 300, the purple strain is present mostly in drug-sensitive infections, whereas the green strain is present in drug-resistant infections. We also see that only three of the strains persist after time 200 and that one of the strains seems to eventually take over the population. The nature of these strains can be predicted before running the simulation by calculating the two eigenvalues from equation 4. Based on equation 10, we also know whether this fittest strain is found mostly in drug-resistant or drug-sensitive infections.

**Figure 3.**
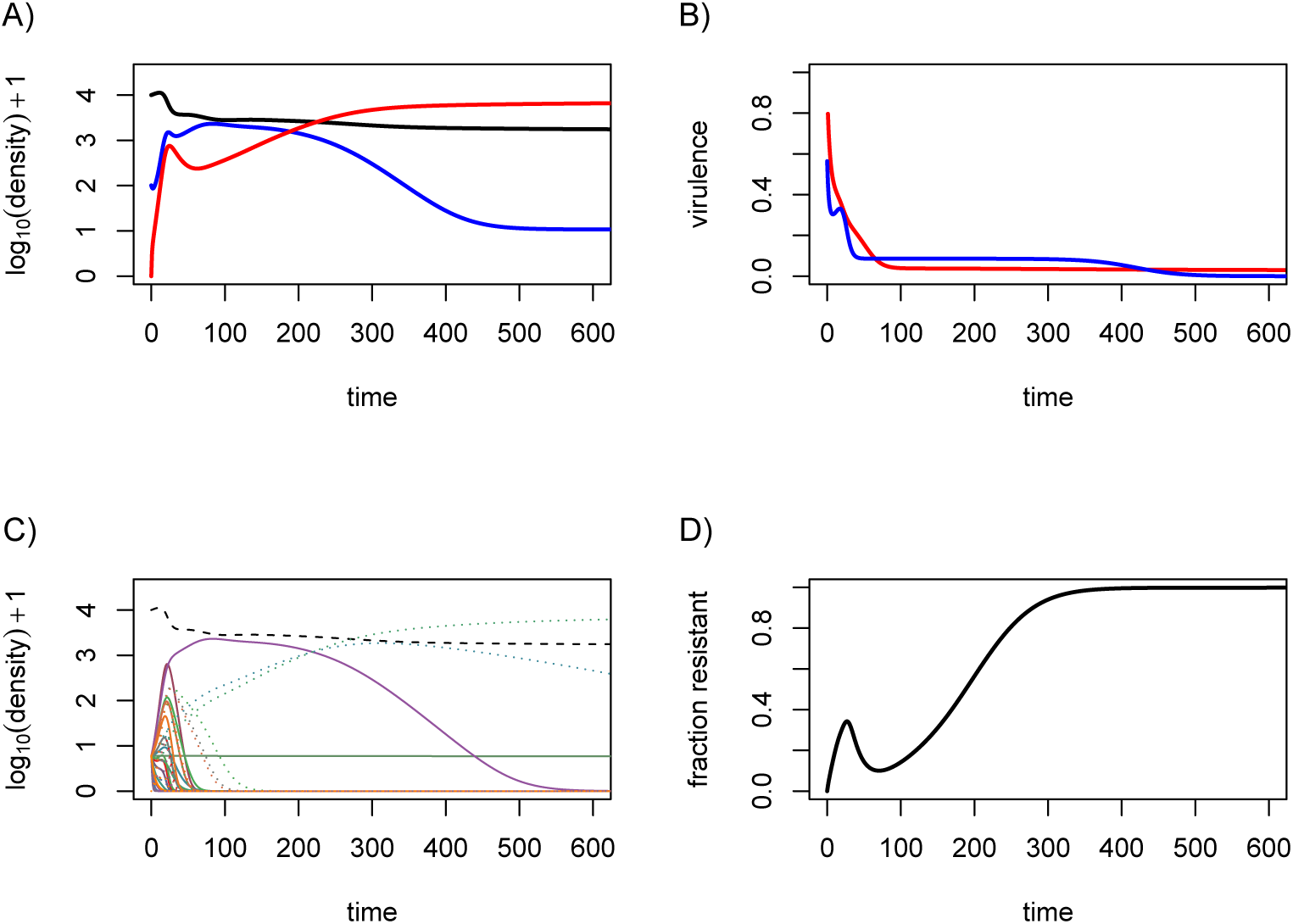
Long term evolutionary dynamics. A) Dynamics of the densities of susceptible hosts (black), drug-susceptible infections (blue) and drug-resistant infections (red). B) Average virulence in the drug-sensitive (blue) and drug-resistant infections (red) for numerical multi-strain simulations. C) Dynamics of the densities of susceptible hosts (dashed black line), of drug-susceptible infections (plain coloured lines) and drug-resistant infections (dotted coloured line). Each colour corresponds to one of the *n* = 20 strains. D) Fraction of drug-resistant infections. Parameter values are the same as in Figure 2 for the multi-strain simulation.

Since the system only contains 20 strains, the presence of drug resistance at equilibrium does not only depend on the model parameter. In Supplementary Figure S2, I show that the presence of drug resistance can be negligible with the same parameter values but with 20 other strains chosen at random using the same distribution.

### Adaptive dynamics

The adaptive dynamics approach can be seen as the opposite of the Price equation approach: the latter is accurate in the short term, whereas the former is accurate in the long term.

The major problem with adaptative dynamics is that it requires making assumptions about trade-off relationships, and we currently lack detailed biological data regarding most of these trade-offs. Generic qualitative trends can still be analysed by, at the same time, results based on critical function analysis show that trade-off relationships with only slight differences can lead to qualitatively different evolutionary outcomes (Kisdi, 2006; Svennungsen and Kisdi, 2009). Therefore, this final subsection should be more interpreted as the illustration of a potential scenario rather than a widespread system behaviour.

### ESS values

Using the adaptive dynamics approach, it is possible to know which of the strains present in the system will eventually persist. Since there is no density-dependent feedback (eigenvalues only depend on 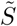 and not on 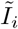 or 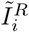). As shown in the methods section, there are two expressions for the dominant eigenvalue (equation 4). Based on earlier work, we know that these two eigenvalues either yield a null virulence, a maximal virulence, or an intermediate evolutionarily stable (ESS) level of virulence depending on the exact shape of the relationship between transmission rate and virulence (van Baalen and Sabelis, 1995).

The ESS eventually reached, provided that parameter values allow for its existence, is the one that maximises its fitness as expressed in equation 4. The first candidate ESS value can readily be obtained using classical approaches if we assume the generic transmission-virulence trade-off from equation 6 with 0 < *p* < 1 and assuming that virulence in drug-resistant infections is proportional to that in drug-sensitive infections 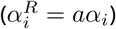:

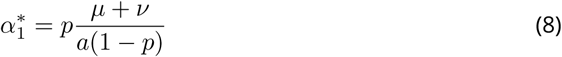

as shown previously, we find that there can only be a finite non-zero ESS if 0 < *p* < 1.

The second candidate ESS value is less trivial as it also relies on the shape of the trade-off between treatment rate and virulence. As shown in the Appendix, we can derive a condition that 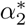 must satisfy, assuming the same transmission virulence trade-off function:

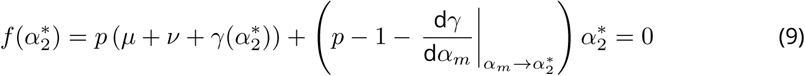

We know from condition S13b that *f* is a decreasing function of 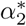. Therefore, increasing the value of a parameter in *f* will require a larger value of *α*\* such that *f*(*α*\*) = 0. For instance, increasing the recovery rate (*ν*) selects for higher virulences, as expected (van Baalen, 1998). We see that increasing the intensity of the treatment rate favours more virulent strains. However, the faster the treatment rate increases with the virulence, i.e. the larger d*γ*/d*α*, the lower the ESS virulence.

### Multiple equilibria

One less common feature of the current system is that, as the virulence of the resident strain evolves via successive invasion/replacement events characteristic of the adaptive dynamics approach, it is theoretically possible for the system to switch between the two steady states. Indeed, based on equation 4, the fittest strain, denoted with a star (*), is the one with the highest value of

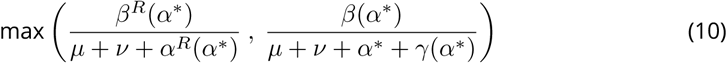

From this equation, we see that without a cost associated with drug resistance, i.e. if *β^R^* ≥ *β* and *α^R^* ≤ *α*, the system will always tend towards drug-resistant infections as soon as the treatment rate is non-zero (and provided that there is sufficient genetic variation).

We can also formulate a qualitative analysis of this simple system and distinguish between two evolutionary scenarios. First, if one type of infection state (drug-resistant or drug-sensitive) always has higher fitness than the other, then virulence will evolve to this single peak. Second, if the nature of the infection state with the highest fitness depends on the virulence value, then the initial virulence may lead the system to evolve to a local maximum. As shown in Figure 4, if the initial virulence is zero then the system will converge to the ESS virulence corresponding to the drug-sensitive equilibrium (dotted yellow line). A large mutation event, e.g. the emergence of a very virulent strain, could move the system to the drug-resistant ESS (plain red line), which is a global maximum. Note also that increasing the scaling parameter between treatment rate and virulence favours less virulent strains in the drug-sensitive equilibrium (dotted blue line). However, the decrease in virulence (i.e. the shift of the fitness peak to the left in Figure 4), also comes with a decrease in parasite fitness, which can make the drug-resistant equilibrium (red plain line) globally stable.

**Figure 4.**
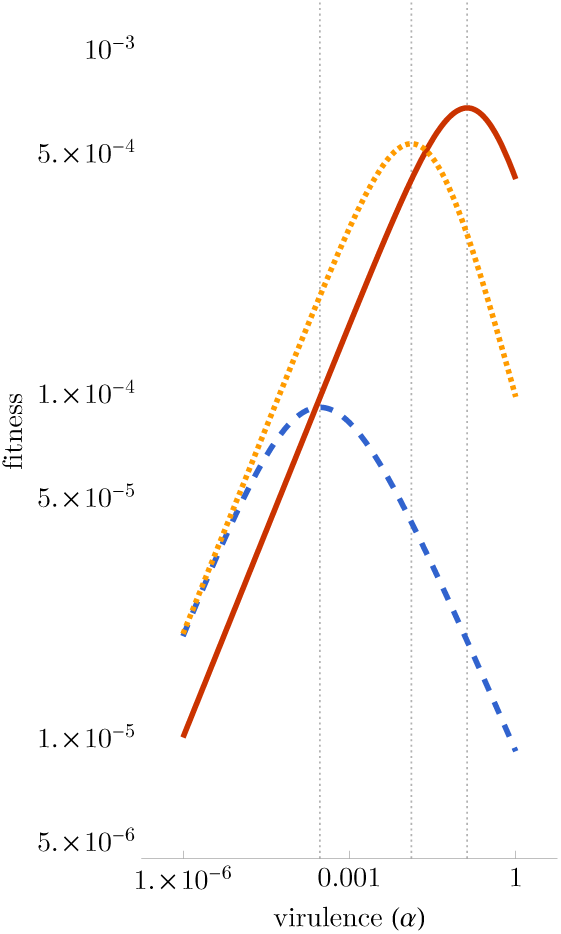
Fitness of drug-resistant (plain red) and drug-sensitive (dotted blue and yellow) infections as a function of virulence. This illustration assumes trade-offs between virulence and transmission rate (*β*(*α*) = β0*α^p^*) and treatment rate and virulence (*γ*(*α*) = *g α^q^*). In the blue dashed case, treatment rate is more intense against virulent infections (*g* = 10) than in the dotted yellow case (*g* = 0.1). Vertical lines show the ESS values. Parameter values are *β*_0_ = 10^−4^, *p* = 0.4, *q* = 0.75, *μ* + *ν* = 0.02, *a* = 0.1 and *b* = 0.5.

From a public health perspective, this means that targeting more virulent infections with higher intensity can be an interesting strategy because it favours less virulent strains. However, a potential risk is that the local ESS with a drug-sensitive infection disappears and that the drug-resistant ESS becomes globally stable. More realistic functional links between treatment rate and virulence could affect these results and even generate more elaborate eco-evolutionary feedbacks, for instance, if the number of infected hosts affects the treatment rate (Dieckmann, 2002; Pharaon and Bauch, 2018).

## Discussion

As attested by the resolution from the United Nations in 2016, anti-microbial drug resistance is a world issue with strong implications for public health policies (Wernli et al., 2017). However, efforts are still required to popularise it as an evolutionary process (Antonovics et al., 2007). This is not a mere semantic issue because it means envisaging antimicrobial resistance as a dynamical process, which directly impacts the design of optimal therapies (AF Read, Day, et al., 2011; Wiesch et al., 2011). Using an epidemiological modelling framework to analyse short-term and long-term evolution, I show that public health policies can generate differences in virulence between drug-resistant and drug-sensitive infections.

First, assuming that more virulent infections are more symptomatic and, hence, more treated, the initial emergence of drug-resistance is expected to occur first in more virulent strains. Second, even if drug-resistance pre-exists in the population, virulence should initially be less counter-selected in the drug-resistant population, leading to drug-resistant infections being less virulent on average. The more the epidemic unfolds, the more this gap can widen or shrink.

Indeed, in the long term, the nature of the strain that takes over the host population results from genetic trade-offs. In this simple model, we find using an adaptive dynamics approach that the strain that takes over the population will generate either mainly drug-sensitive or drug-resistant depending on the nature of the fitness costs associated with drug resistance. These two outcomes correspond to two different ESS. In the case where drug-sensitive infections dominate, we find a positive correlation between treatment rate and virulence selects for less virulent strains. However, depending on the parameter values, the risk is that targeting more virulent infection (i.e. increasing the correlation) may lead the ESS associated with drug-resistance to become globally stable, thereby rendering the new ESS value independent from the treatment rate.

### Biological implications

One direct consequence of these results has to do with the estimation of infection virulence, especially in an outbreak situation. Indeed, in addition to controlling for host differences that can affect virulence (since it is a shared trait between the host and the parasite), the drug-resistant nature of the infection should also be taken into account since, initially, virulence is expected to be higher in drug-resistant infections. The advantage of drug-resistance compared to other biases, such as reporting rate, is that it can be assessed *a posteriori* on clinical samples. The downside is that there can be a direct link between drug resistance and virulence. Ideally, one would want to also assess parasite virulence *in vitro*, but this is much less straightforward than drug resistance.

There are several ways to test the predictions of this study. One possibility could be to use an experimental system where treatment can be applied to hosts based on the symptoms they exhibit. One difficulty is that it requires to have hosts large enough to detect the symptoms and apply treatment without affecting the rest of the population. Another difficulty is that it imposes to be able to generate real epidemics. Fungal or bacterial parasites of plants could provide the ideal system. Another possibility would be to analyse existing clinical or agronomical data. A potential parasite could be HIV infections because drug resistance is well characterised (Beerenwinkel et al., 2003) and virulence can be estimated through set-point virus load (Fraser, Lythgoe, et al., 2014). However, a major difficulty is that to test the model, we would either need to know the nature of the fitness cost potentially associated with drug-resistant mutations. A possibility could be to use algorithms that can partly predict virus load based on the virus sequence (Hinkley et al., 2011).

### Simplifying assumptions

Several simplifying assumptions were made to analyse the model. One of them is that drug resistance is a binary trait when virulence is a continuous one. Modelling drug-resistance as a continuous trait is possible. However, from a modelling point of view, unless we have detailed data to calibrate mutation kernels, it would be impossible to distinguish strain-level variations in the level of virulence from variations in the level of resistance. As a consequence, the feedback between the ecology (treatment policies) and trait evolution (drug resistance) would disappear. Conversely, it would be possible to model virulence as a discrete trait. This would have the advantage that by modelling the system with (at least) two loci, one governing the drug resistance phenotype and another governing the virulence phenotype, we could draw parallels with and perhaps use data on virulence factors. Multi-locus drug resistance mathematical model already exist that can readily tackle this question (Day and Gandon, 2012). Unfortunately, this binary structure is poorly adapted to capture quantitative variations in virulence, which have been shown to lead to trade-off relationships with other traits such as transmission rate (Anderson and May, 1982; de Roode et al., 2008; Doumayrou et al., 2012; Dwyer et al., 1990; Fraser, Hollingsworth, et al., 2007; Raberg, 2012; Williams et al., 2014). In the end, the main results would most likely depend on the genetic linkage between the two loci and on the assumption regarding the fitness effect of the alleles at the virulence locus (Day and Gandon, 2012).

The model can be extended in several ways. For instance, since demography has no impact, the time scale is flexible, which means the model will exhibit similar dynamics for short infections with a high transmission rate and chronic infections with low transmission rates. Another oversimplifying aspect of this model resides in its inability to discriminate between the rate at which an infection is treated (*γ_i_*) and the probability of treatment failure (*ρ_i_*). One possibility to tackle both limitations simultaneously would be to explicitly model recovered individuals and assume that immune memory is not lifelong. Another extension could be to model symptoms explicitly. However, the problem with splitting the I compartment between a pre-symptomatic and a post-symptomatic stage is that, at least with the Price equation formalism, this would multiply by 2 the number of equations required to follow trait-dynamics, making the short-term evolution results less easy to interpret.

Our formalism has the advantage to allow for a continuum of strains with varying values of virulence and transmission rate. Regarding the latter parameter, one simplifying assumption of the model is that it is defined as a function of virulence. In reality, the realised transmission rate will be distributed around this expected value. As illustrated in Supplementary Figure S3, I also allowed some degree of noise on this transmission rate. In general, the results are qualitatively robust to this noise and the main variable that is affected is the fraction of drug-resistant infections (Supplementary Figure S4). The model also assumes a saturating transmission-virulence trade-off relationship, which means there exists an intermediate optimal level of virulence (Alizon and Michalakis, 2015; van Baalen and Sabelis, 1995). Without it, parasites evolve to avirulence but they still do so more rapidly in the drug-sensitive than in the drug-resistant infections.

### Short *vs*. long-term predictions

The qualitative predictions from the Price equation appear to be correct initially but there is an increasing mismatch for some traits as the epidemic unfolds. This is expected because the variances and covariances are constant parameters in the simulations. However, as the epidemic unfolds, the relative frequency of each of the 20 strains varies over time (as illustrated by Figure 2A) therefore altering the variances and covariance. To take an extreme example, if a single strain dominates, the variances will be 0. Because the formalism used here does not update the covariance matrix, it is bound to eventually be unable to predict the dynamics accurately.

For the adaptive dynamics, we have the opposite issue. On the short term, the variations in virulence or on drug-resistance only depend on the genetic diversity and may be at odds with the predicted ESS value. It is only on the long term that the system will converge towards the evolutionary equilibria predicted by the model Analysing the same system with the Price equation and the adaptive dynamics gives us insight into the early and late evolutionary dynamics without resorting to numerical simulations. Gaining more insights into the system will require more assumptions regarding the underlying correlations between traits. In particular, it would be interesting to allow for compensatory mutations to mitigate the effect of drug resistance costs. Furthermore, modelling explicitly within-host dynamics, and explicitly tracking the competition between drug-sensitive and drug-resistant variants of a given strain through nested models (Mideo et al., 2008) could be a way to study the interaction between levels of adaptation.

## Supporting information

R script to generate the figures

## Supplementary material

The `R` script used to generate the figures is available as a supplementary .txt file online (DOI: 10.1101/2020.02.29.970905).

## Acknowledgements

I thank Mircea T Sofonea and two anonymous reviewers for their comments. This work was supported by the European Research Council (ERC) under the European Union’s Horizon 2020 research and innovation program (EVOLPROOF, grant agreement No 648963). Further support was provided by the CNRS and the IRD. Version 3 of this preprint has been peer-reviewed and recommended by Peer Community In Evolutionary Biology (https://doi.org/10.24072/pci.evolbiol.100113).

## S1 Supplementary methods

### S1.1. Price equation calculations

Following the Price equation formalism introduced by Day and Proulx (2004), we take the derivative with respect to time of *p_i_ = I_i_/I_T_*, we get

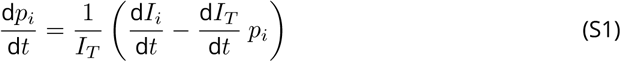

From equations 1b and 2a, we get

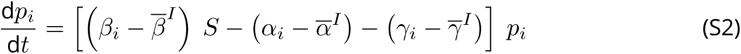

Let us look at the evolution of the average value of a given trait *x*. If we assume that the trait of a genotype *i* does not value with time, we have:

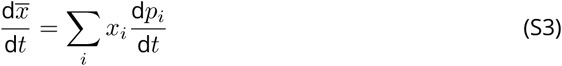

From equation S2, we have:

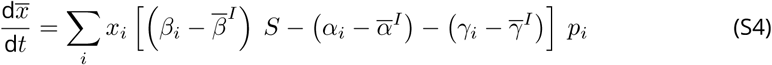

This can also be written as

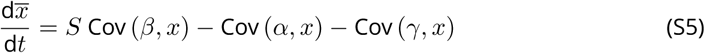

Similarly, we can define the proportion of genotype *i* in the drug-resistant compartment as 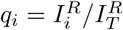 and derive it with respect to time to get:

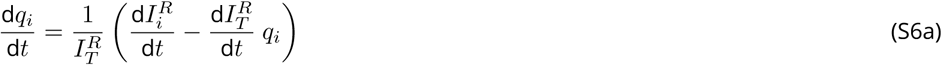

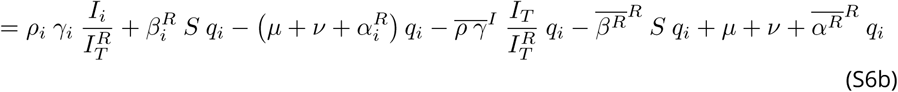

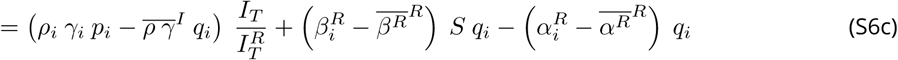

For a given trait *y* in the drug-resistant compartment, we have

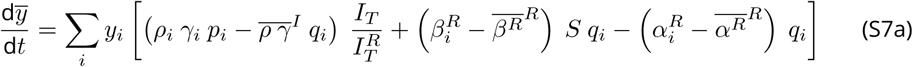

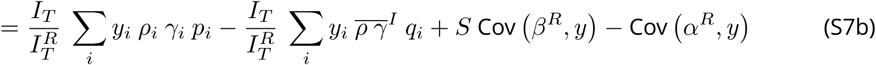

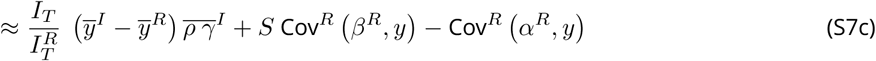

The assumption made to reach the last step is that 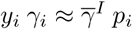.

As before, for the main trait of interest (virulence) and with the same assumptions as before we get:

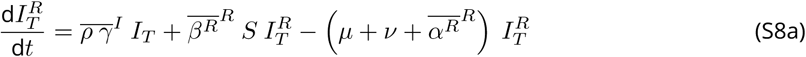

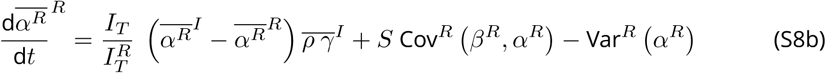

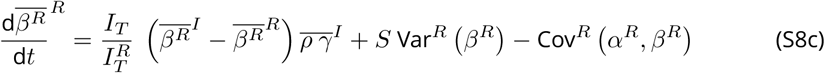

### S1.2. System equilibria

The detailed equilibria of the main equation system are the following:

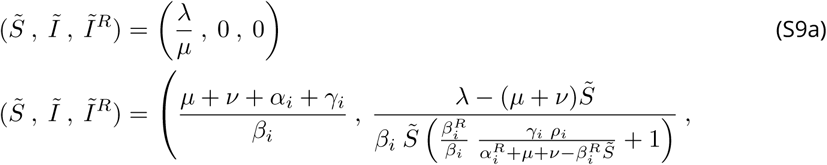

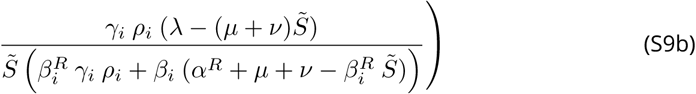

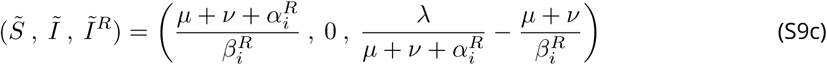

### S1.3. Adaptive dynamics approach

#### Deriving the invasion fitness

We now adopt an adaptive dynamics approach to study long term evolution based on the functional relationship between treatment rate and virulence (*γ(α)*). For generality, we also initially denote mutation rate as a function of virulence (*ρ*(*α*)). Finally, we assume a transmission virulence trade-off (*β*(*α*)) otherwise virulence will not be adaptive and always be selected against.

Let us rewrite the ODE system:

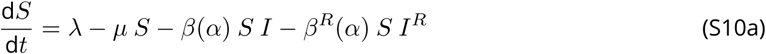

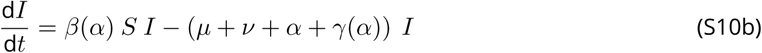

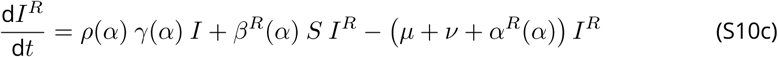

Following the next-generation matrix theorem (Diekmann et al., 1990; Driessche and Watmough, 2002; Hurford et al., 2010), the Jacobian matrix of the system (*J*) can be decomposed into a ‘birth’ (*F*) and a ‘death’ (*V*) matrix:

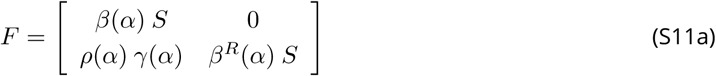

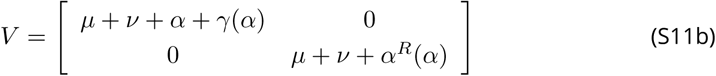

The eigenvalues of *F.V*^−1^ are

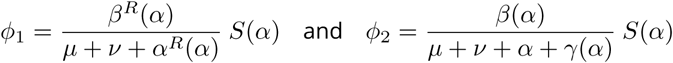

Following the adaptive dynamics framework, we then assume that system S10 is at en epidemiological equilibrium where the parasite persists (see equations S9). This strain is referred to as the ‘resident’ strain and its trait of interest, virulence, is denoted *α_r_*. We then assume that another strain emerges through mutation that has a slightly different trait value *α_m_*. Key assumptions are that the mutant density is rare compared to that of the resident and that its trait value is close to that of the resident.

Note that if we allow for ‘reversions’ of drug-resistant infections, the eigenvalues can still be derived but their expression is less clear. We can also see that any link between treatment failure probability and virulence does not matter on the long run.

A perturbation analysis of the system for a rare mutant where the density of susceptible is set by the endemic equilibrium of the resident strain therefore leads to the following eigenvalues

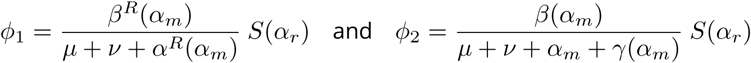

In this simple model, the ESS corresponds to the strategy that maximises these invasion fitnesses (Dieckmann, 2002). Since the resident trait only affects the density of susceptible hosts, the fitness optimum is an absolute maximum and not a relative maximum.

#### Evolutionary singular strategies

The first eigenvalue *φ*_1_ is identical to that of SI systems studied in details in earlier studies. Depending on the concavity of the transmission-virulence trade-off curve, there can be an ESS with an intermediate level of virulence. More precisely, as shown by van Baalen and Sabelis (1995), there is an ESS if there exists a virulence *α*\* that satisfied the two conditions

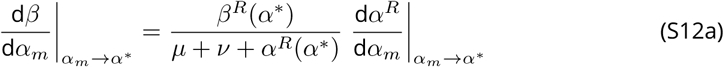

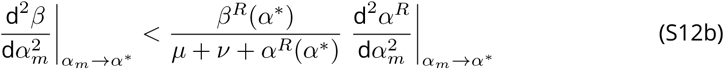

They also show that there is an elegant graphical interpretation to this condition, which is that if there is an ESS, then in *α*\* the tangent of the parametric curve 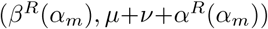 also passes through (*μ* + *ν*, 0). Note that treatment rate (*γ*) has no effect on this ESS.

The second eigenvalue is very similar to the first and, again using the work from van Baalen and Sabelis (1995), it can be shown that the second ESS satisfies the following conditions:

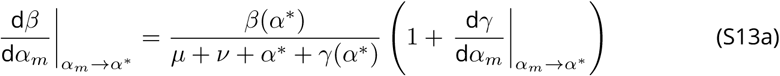

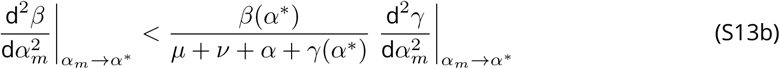

Assuming that conditions S13 are satisfied, we can study the effect of variations in treatment rate on the ESS value, which is found by solving equation S13a. This condition can also be written as

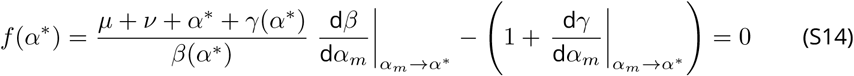

We know from condition S13b that *f* is a decreasing function of *α*\*. Therefore, increasing the value of parameter in *f* will require a larger value of *α*\* such that *f*(*α*\*) = 0. For instance, increasing the recovery rate (*ν*) selects for higher virulences, as expected (van Baalen, 1998). We see that increasing the intensity of the treatment rate favours more virulent strains. However, the faster the treatment rate increases with the virulence, i.e. the larger *γ*’(*α*), the lower the ESS virulence.

Notice that with the transmission-virulence trade-off function assumed in the main text, we have, for 0 < *p* < 1,

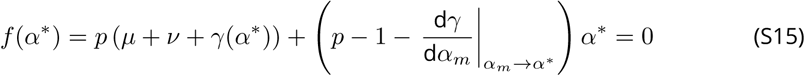

## S2 Simulation R code

This appendix contains the `R` code used for the simulations and generating the figures.

## S3

**Figure S1.**
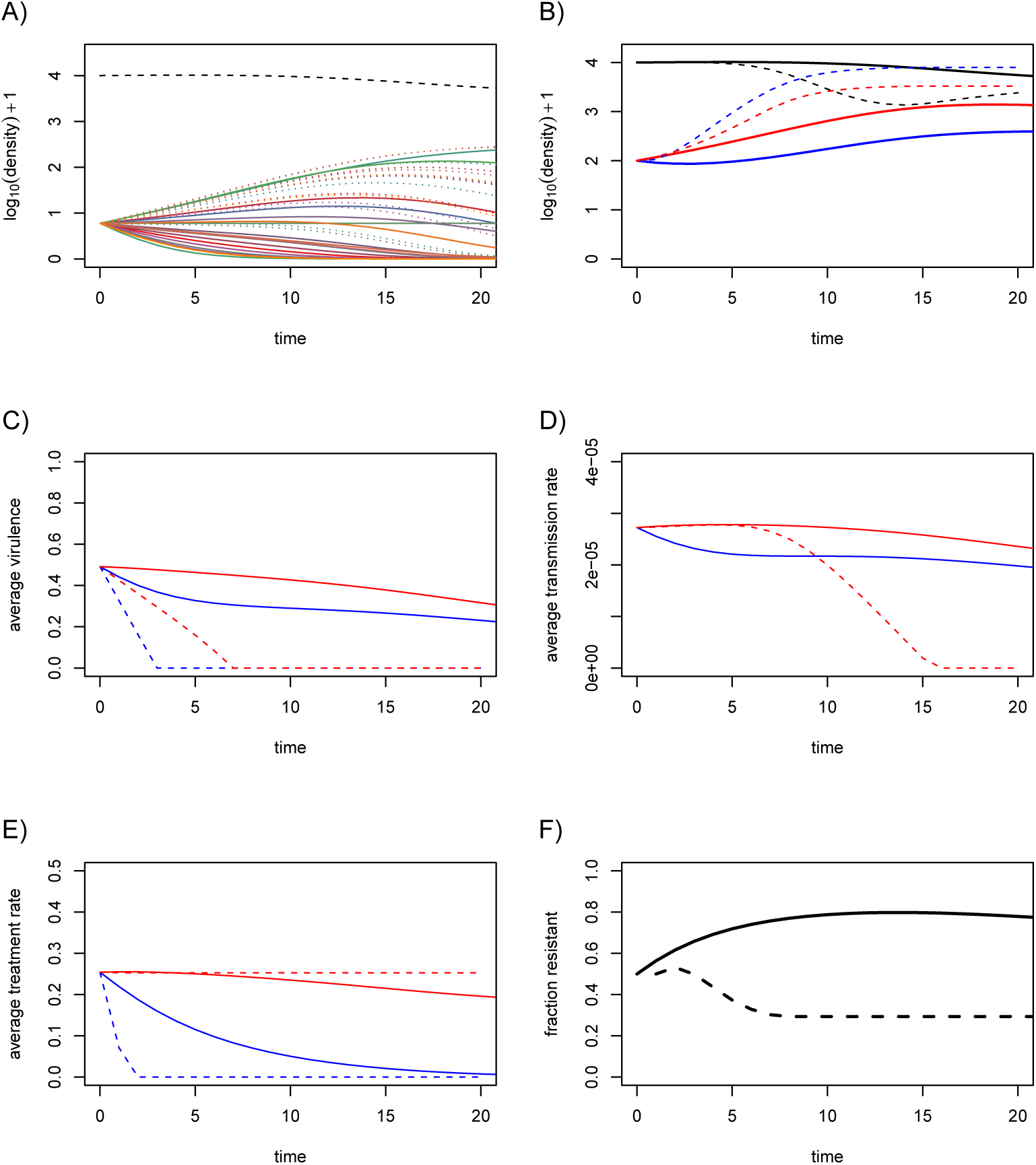
Short-term dynamics with pre-existing drug resistance. Virulence in the drug-resistant compartment still decreases more slowly than in the drug-sensitive compartment if we assume that the initial density of the two types of infections is equal (therefore that drug resistance is not only generated by treatment failure initially). 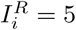 and other parameter values are identical to that in the main text.

**Figure S2.**
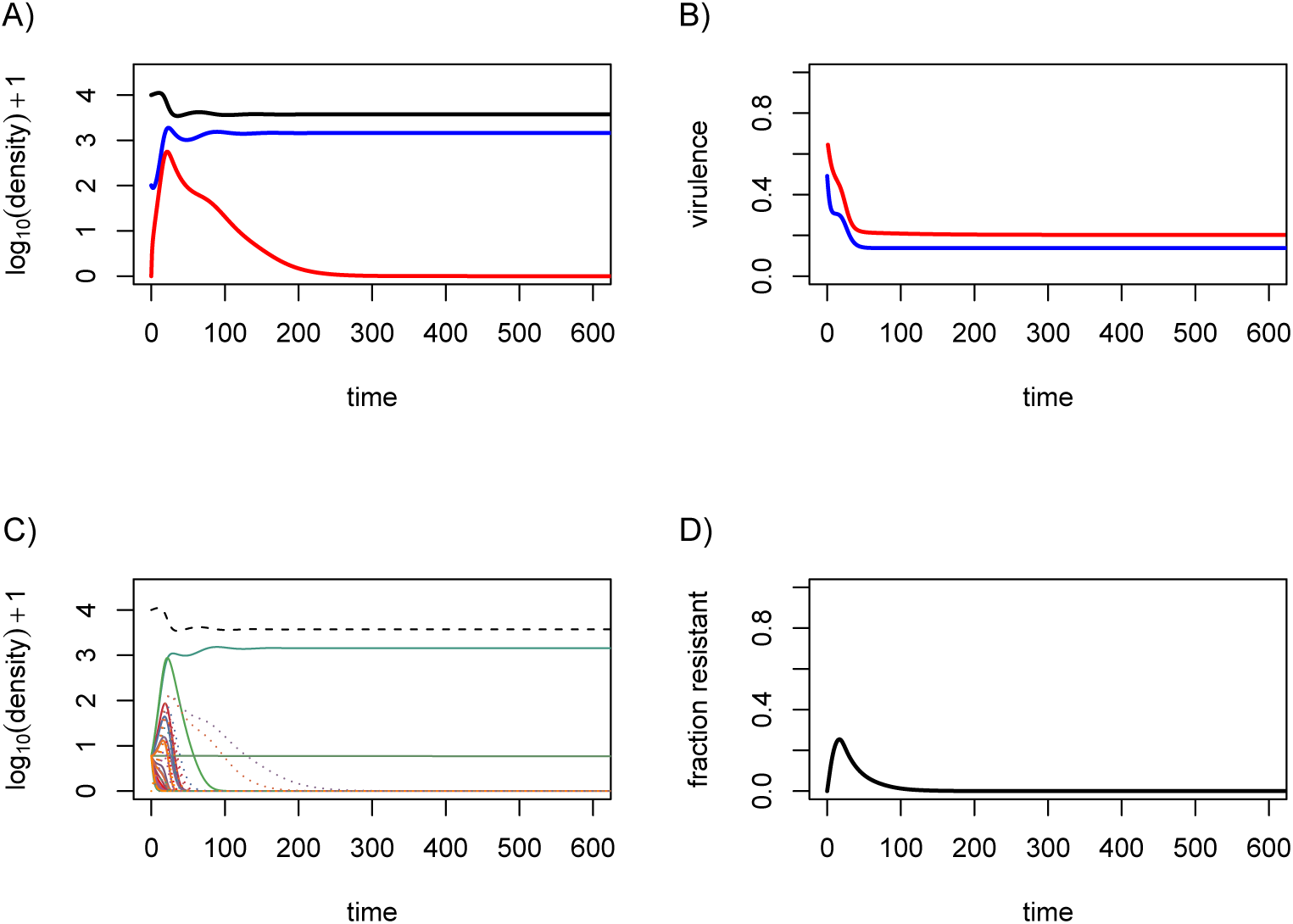
Long-term evolutionary dynamics. Even with the same parameter values as in the main text, drug-sensitive infections can eventually dominate the system. This is because we only have a limited number of strains in the simulations. Parameter values are the same as in the main text but the *n* = 20 strains are different (although drawn using the same covariance matrix as in the main text).

**Figure S3.**
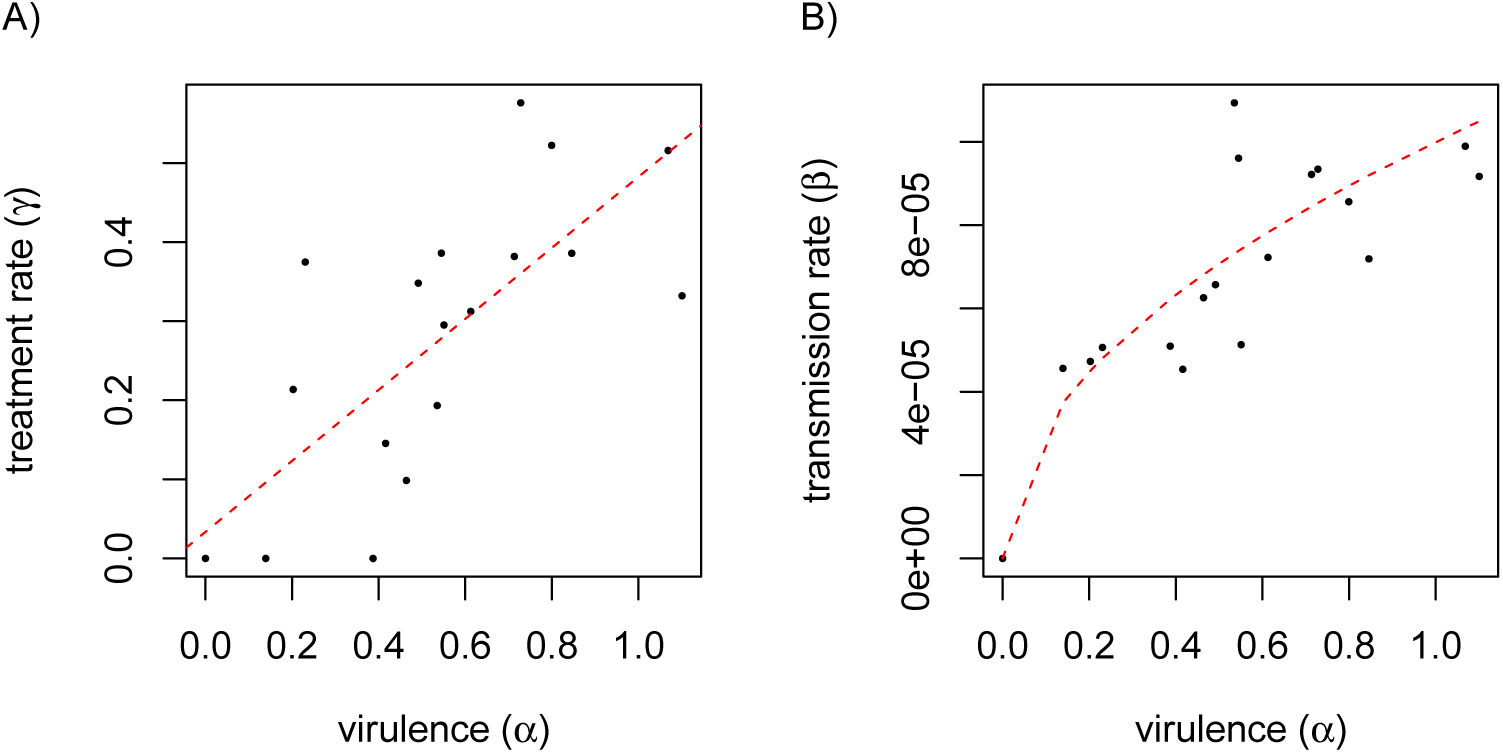
Trade-off relationships between model parameters. A) Positive covariance between virulence and treatment rate (values are drawn from a multivariate distribution). B) Trade-off relationship assumed between transmission rate and virulence and points obtained when by adding noise to the transmission rates (normal distribution with mean 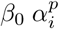 and variance 0.2). Dashed lines show the results of a linear model fit between *α* and *γ* in panel A, and the theoretical trade-off relationship from equation 5 for panel B.

**Figure S4.**
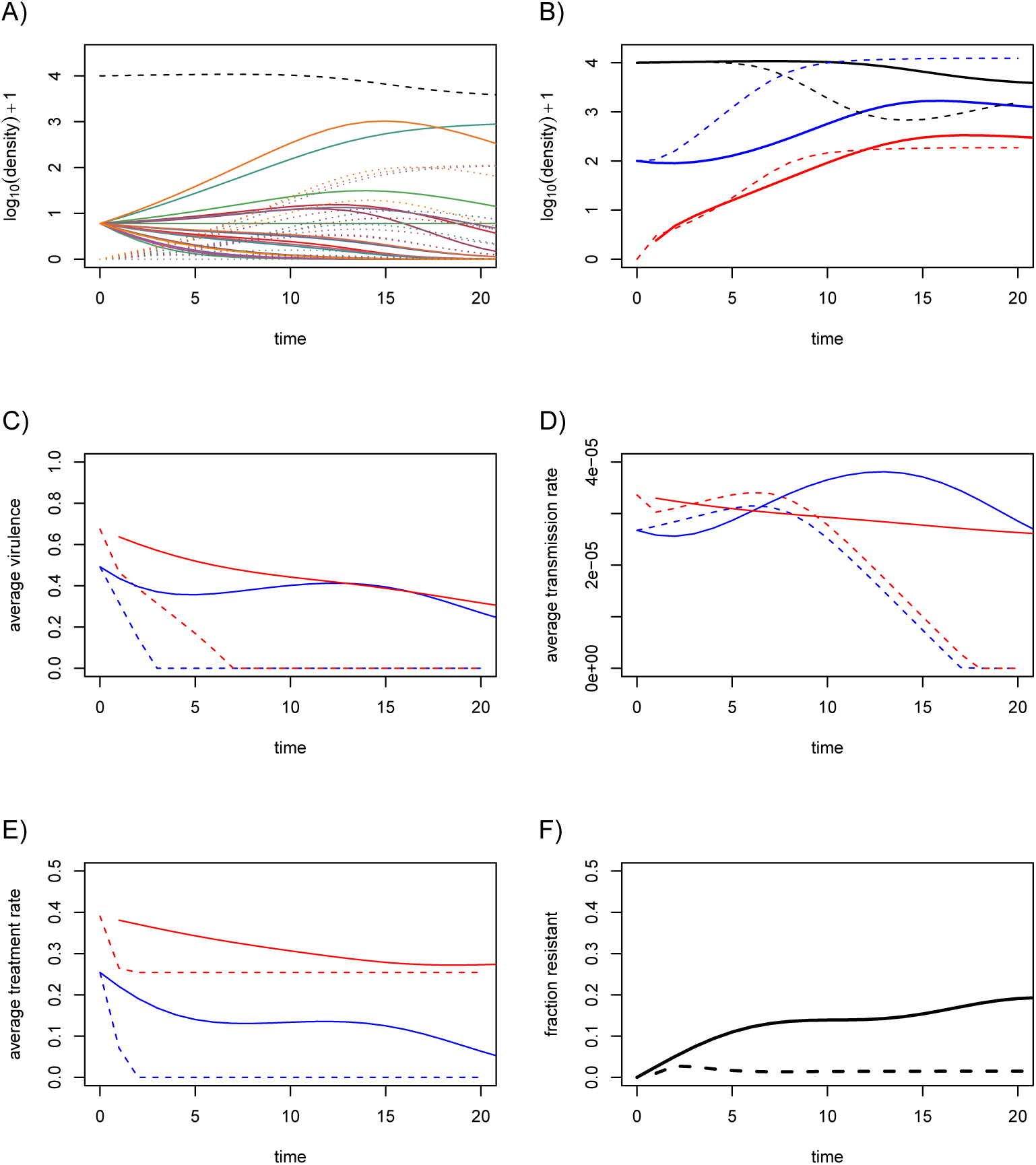
Short-term evolutionary dynamics assuming noise in transmission rates. Figure captions are identical to that in the main text. Parameter values are identical to that in the main text, except for the noise, which is generated by a Gaussian distribution centred around the transmission value predicted by the trade-off shown in Figure S3B and with relative standard deviation 20%.

